# Identification of molecular and clinical ALS subgroups based on TDP-43 loss of function molecular markers from population-based patient-derived iPS motor neurons

**DOI:** 10.64898/2025.12.28.696512

**Authors:** Trinity Cheng, Shaili Tripathi, Yike Guo, Parimala Vedula, Ryder Li, Michael Potanin, Nidhi Soley, April Yujie Yan, Ishan Vatsaraj, Carl Harris, Joseph L. Greenstein, Casey Overby Taylor, Alyssa N. Coyne, Jeffrey D. Rothstein

## Abstract

**Background:** Amyotrophic lateral sclerosis (ALS) is a uniformly fatal neurodegenerative disease characterized by progressive cortical and spinal motor neuron loss, with most patients surviving only 2–5 years post-diagnosis. While approximately 10% of cases are familial (fALS), the remaining 90% are sporadic (sALS) with unknown genetic drivers. Importantly, clinical presentations are heterogeneous in both sporadic and familial ALS, underscoring the complexity of the disease. A pathological hallmark of ALS is the mislocalization of RNA-binding protein TDP-43 from the nucleus to the cytoplasm. This mislocalization produces both loss of function consequences, such as widespread RNA processing and splicing defects, as well as potential toxic gain of function effects associated with cytoplasmic aggregation.

**Results:** In this study, we used RT-PCR data from induced pluripotent stem cell-derived motor neurons derived from 180 sALS and C9orf72 fALS patients from the Answer ALS collection to identify biological subgroups based on TDP-43 loss-of-function signatures. Spectral embedding revealed four distinct molecular clusters, including one subgroup genetically similar to controls and another with the most dysregulated mRNA expression, suggesting differing disease severity. Linear mixed models were then used to assess the longitudinal trajectory of over 90 clinical measures, and the between-cluster interaction effects were evaluated.

**Conclusions:** 36 clinical outcomes showed significant differences across clusters, supporting the presence of biologically and clinically distinct ALS subtypes based on the TDP-43 associated pathogenic cascade. These findings demonstrate a critical role of RNA profiling in uncovering biologically meaningful subtypes of ALS, potentially allowing for more precise prognostic tools and the development of future personalized therapeutic approaches.

## Introduction

Amyotrophic lateral sclerosis (ALS) is a fatal and clinically heterogeneous neurodegenerative disease that primarily affects upper and lower motor neurons. More than 50 ALS-associated genes have been identified, including *C9orf72, SOD1*, and *FUS*, which contribute to individual variations in disease onset, symptomatology, and rates of progression^12^. This heterogeneity complicates efforts to provide accurate assessments of individual risk and prognosis, design targeted therapies, and optimize treatment timing. Although approximately 10% of cases are familial (fALS), the vast majority (90%) are sporadic (sALS), with no family history or clearly defined genetic causes. Stratifying patients into biologically meaningful subgroups could improve clinical care by enabling tailored treatment strategies, more accurate prognostic discussions, and perhaps most importantly, personalized therapeutic interventions. Such stratification could also strengthen clinical trial design by reducing heterogeneity and improving the likelihood of detecting treatment effects.

Efforts to categorize ALS patients include criteria such as family history, clinical milestones, neurophysiological assessments, and various serum biofluids such as neurofilament light chain^3–6^. While useful in deepening our understanding of the disease, these classifications often reflect symptomatic distinctions rather than true biological or molecular subgroup differences, limiting their utility and applicability in a clinical setting, especially with regards to therapeutic development or optimization. For example, semi-supervised machine learning models were able to identify clusters corresponding to the six clinical subtypes defined by the Chiò classification system^4^. However, the Chio system primarily divides patients by site of disease onset and has limited therapeutic implications, as it cannot be used to recommend treatments or predict prognoses. Even recent studies that utilize molecular and biochemical data rather than solely clinical data have experienced difficulty identifying useful subgroups. In 2024, a comprehensive multi-omics characterization of the prefrontal cortex (PFC) was conducted for 51 ALS patients, confirming that molecular subclusters drive heterogeneity in the PFC^7^. However, the differences in molecular mechanisms that were identified were primarily sex-specific, a division that is interesting but not clinically meaningful for treatment purposes. Some studies have utilized central nervous system autopsy tissue transcriptomics, but suffer from imprecise molecular pathways and the serious artifacts of postmortem or agonal-related events^8,9^.

The Answer ALS Research Project, the most comprehensive ALS multi-omics dataset to-date, was developed to drive the identification of clinical-molecular-biochemical subtypes of ALS at a population level^10^. Deep longitudinal clinical data as well as comprehensive multi-omic analytics from induced pluripotent stem cell (iPSC) lines are available for more than 1000 ALS and control patients, resulting in around six billion data points per patient awaiting deeper analysis. This study focuses on a cohort of 180 ALS patients from the Answer ALS dataset, for whom additional targeted qRT-PCR data from iPSC-derived motor neurons (iPSNs) is available^11,12^. ALS is a TAR DNA-binding protein 43 (TDP-43) proteinopathy, with nuclear depletion of TDP-43 compromising its nuclear function in 97% of both fALS and sALS patients. TDP-43 dysfunction contributes to widespread RNA processing defects, protein dysregulation, and neurodegeneration across multiple cellular pathways^13^. A recent iPSN dataset evaluated multiple TDP-43 mRNA targets to define TDP-43 loss of function signatures in a large cohort of sALS patient cells. We used these datasets in attempt to advance ALS subgroup identification through a machine learning approach that integrates traditional clinical and phenotypic data with multi-omics analytics curated to cover the spectrum of significant ALS hallmarks, particularly TDP-43 loss of function. Most prior ALS research has centered on isolated genomic or phenotypic datasets only, often without a focus on ALS-specific pathways. This data largely derives from the living nervous system iPS cells from a large population of individual patients, serving as a biopsy-like surrogate. Importantly, TDP-43 and nuclear pore defects seen in Answer ALS iPS neurons mirror those in the patients’ own autopsy tissue, confirming that this model faithfully reproduces these disease pathways (cite Rothstein/Coyne Nat Commun 2025).

By utilizing this combination of comprehensive, novel data and powerful data science tools, we aim to identify new, biologically meaningful disease subgroups that focus on the neurological underpinnings of ALS, and uncover subgroup-specific mechanisms that may drive disease progression.

The primary objective of this study was to develop a machine learning (ML)-based prognostic model that leverages these transcriptomic profiles to stratify ALS patients into biologically meaningful subgroups. Such subgroups, ideally representing distinct disease trajectories or biological pathways, would allow for improved clinical decision-making and tailored therapeutic development. Specifically, the goals were to: (1) identify molecular subgroups of sporadic ALS patients based on gene expression and splicing signatures associated with TDP-43 loss of function using iPSC-derived spinal motor neuron transcriptomic data from the Answer ALS program; (2) characterize molecular differences among these subgroups by comparing expression and splicing changes in known TDP-43-regulated genes using non-parametric statistical analyses; and (3) examine clinical differences across subgroups by modeling longitudinal outcomes such as ALS Functional Rating Scale (ALSFRS) scores, motor strength, and respiratory function with linear mixed-effects models (LMMs).

## Methods

### Dataset Overview

This study draws on data from Answer ALS, program (www.neuromine.org), which includes clinical, demographic, and multi-omics profiles from over 1,000 patients5. The dataset provides clinical demographics (age, sex), indices of disease progression (ALSFRS scores), and clinical measurements (respiratory function, fine motor activity, neurological assessments), as well as multi-modal biological data from induced pluripotent stem cell (iPSC) lines derived from each participant.

From Answer ALS, 180 iPSC lines were differentiated into spinal motor neurons (iPSNs), and profiled for gene expression and exon-level splicing^6^. This qRT-PCR based data set was specifically designed to assess molecular signatures associated with TDP-43 loss of function, comprising of 6 gene expression changes (ELAVL3, PFKP, RCAN1, SELPLG, STMN2, and UNC13A), 12 TDP-43 loss of function related cryptic exon inclusion splicing events (ACTL6B, ARHGAP32, CAMK2B, CDK7, DNM1, HDGFL2, MYO18A, NUP188, POLDIP3, SYT7, STMN2, and UNC13A), and 2 negative controls (ACTIN and the nucleoporin POM121) not known to be regulated by TDP-43. qRT-PCR was performed at three time points (Days 32, 46, and 60) during differentiation to capture the evolution of molecular changes over time. Patients with fALS were filtered out, resulting in 141 remaining sALS patients. The generation and characterization of these neurons, TDP-43 profiling, and pathogenic cascades has been previously described^12,14,15^. All individual iPS lines transcriptomic data and clinical data has been previously published^10,12^.

Demographic data (e.g., sex, age, race, and family history) and longitudinal clinical measurements of ALS disease indicators (e.g., ALS Functional Rating Scale scores, reflex testing, and handheld dynamometry readings) were obtained for these 141 patients from the publicly available Answer ALS/Neuromine database (www.Neuromine.org)^10^. The number of data measurements per patient varied, as did the period of time elapsed between measurements.

### Clustering iPSN Data to Identify ALS Patient Subgroups

iPSN data was examined using two-tailed t-tests to visualize differences in gene expression between the ALS and control patients at each of the three time points. The same tests were conducted between male and female ALS patients.

Prior to clustering, five dimensionality reduction methods were evaluated to remove noise and highlight dominant patterns of variation: Principal Component Analysis (PCA), t-distributed Stochastic Neighbor Embedding (t-SNE), Uniform Manifold Approximation and Projection (UMAP), Spectral Embedding, and Pairwise Controlled Manifold Approximation Projection (PaCMAP)^16–20^. All five methods were implemented in Python, with PCA, t-SNE, and spectral embedding from the scikit-learn Python library. For each method, the Davies-Bouldin (DB) index and Calinski-Harabasz (CH) index were calculated to assess clustering quality based on compactness and separation^21,22^. After dimensionality reduction, K-means clustering was applied, with the optimal number of clusters determined based on within-cluster sum of squares (WCSS). As methods like t-SNE and spectral embedding are sensitive to parameter choices, multiple configurations were tested to optimize performance for each method before comparing.

### Analyzing Molecular Differences Between ALS Subgroups

To explore the molecular distinctions among the identified ALS subgroups, each patient was assigned their new cluster labels, and differential gene expression analysis was conducted across these clusters. The Kruskal-Wallis test, a non-parametric method for comparing non-normally distributed data, was used to assess whether there were significant differences in gene expression levels across the clusters^23^. For any genes found to be significantly different, post-hoc pairwise comparisons using the Wilcoxon Rank-Sum test were conducted to further identify specific inter-cluster differences. The distributions of gene expression across clusters were visualized (Fig. 2), providing insight into the potential mechanisms driving disease heterogeneity.

Subgroup discrimination analyses were performed to further assess whether differences in TDP-43– associated cryptic exon (CE) expression were sufficient to distinguish molecular subgroups. For each subgroup, one-vs-rest classification was conducted using logistic regression with L1 regularization, which promotes sparse feature selection by retaining only features that contribute to subgroup discrimination. Model performance was evaluated using five-fold stratified cross-validation, summarized by the area under the receiver operating characteristic curve (AUC). To assess feature sufficiency, classifiers were re-fit using increasing numbers of top-ranked features, allowing determination of the minimal feature subset required to achieve peak discriminative performance. CE features with non-zero coefficients were considered selected, and coefficient sign was used to indicate whether increased CE expression was associated with increased or decreased probability of subgroup membership.

### Identifying Clinical Differences Between ALS Subgroups

From 44 categories of clinical data, data in administrative or protocol-related categories were excluded from analysis. Additionally, data which was highly variable and consisting of free-text, such as physician notes, was also excluded. Hand Held Dynamometry (HHD) scores, a measure of strength, were averaged across three trials for an average score per muscle^24^. Vital capacity scores for slow vital capacity and forced vital capacity were also averaged across three trials^25^. LMMs were independently fitted for each of the remaining clinical outcomes using the statsmodels Python library^26^. Such models were selected due to their suitability for analyzing longitudinal clinical data, as they account for both fixed and random effects^27^. In this analysis, the fixed effects, representing variables of interest consistent across individuals, were time (days since first measurement), cluster assignment, and the time-cluster interaction. Patient IDs were included as a random effect to model within-subject correlations and adjust for individual differences in disease progression or baseline function. Between-cluster interaction effects from the LMMs were used to compare rates of clinical progression across clusters, capturing variation in symptom trajectories over time.

## Results

### Cohort Characteristics

Patient demographics and clinical characteristics were summarized to confirm comparability across the identified ALS subgroups. Age, sex, and race distributions were examined and reported in Table 1. The mean age of the final cohort was 58.8 years with a standard deviation of 10.5. 67.4% of patients were male, and 31.2% were female, with the vast majority of patients being white. There were no significant differences between ages of the various subgroups. The male:female distribution was similar for each subgroup, with male predominance (well known for ALS population incidence), except for the C9orf72 subgroup.

**Table 1.** Demographics of the 141 ALS patients and control subjects studied.

| Characteristic | Control<br>(N=33) | ALS<br>Overall<br>(N=141) | sALS-1<br>(n=51) | sALS-2<br>(n=60) | sALS-3<br>(n=18) | C9orf72<br>(n=12) |
| --- | --- | --- | --- | --- | --- | --- |
| Age, years (mean ± SD) |  |  |  |  |  |  |
| Age | 57.6 ± 15.9 | 58.8 ± 10.5 | 57.7 ± 11.9 | 59.2 ± 10.8 | 62.7 ± 7.4 | 59.0 ± 5.0 |
| Sex, count (%) |  |  |  |  |  |  |
| Male | 14 (42.4%) | 95 (67.4%) | 33 (64.7%) | 42 (70.0%) | 13 (72.2%) | 7 (58.3%) |
| Female | 19 (57.6%) | 44 (31.2%) | 18 (35.3%) | 16 (26.7%) | 5 (27.8%) | 5 (41.7%) |
| Unknown | 0 (0.0%) | 2 (1.4%) | 0 (0.0%) | 2 (3.3%) | 0 (0.0%) | 0 (0.0%) |
| Race, count (%) |  |  |  |  |  |  |
| White | 28 (84.8%) | 130 (92.2%) | 49 (96.1%) | 52 (89.2%) | 17 (94.4%) | 12 (92.3%) |
| Black / African American | 0 (0.0%) | 9 (6.4%) | 2 (3.9%) | 6 (9.8%) | 1 (5.6%) | 0 (0.0%) |
| Asian | 0 (0.0%) | 0 (0.0%) | 0 (0.0%) | 0 (0.0%) | 0 (0.0%) | 0 (0.0%) |
| Native Hawaiian / Pacific Islander | 0 (0.0%) | 0 (0.0%) | 0 (0.0%) | 0 (0.0%) | 0 (0.0%) | 0 (0.0%) |
| American Indian / Alaska Native | 0 (0.0%) | 2 (1.4%) | 0 (0.0%) | 1 (1.6%) | 0 (0.0%) | 1 (7.7%) |
| Hispanic | 1 (3.0%) | 0 (0.0%) | 0 (0.0%) | 0 (0.0%) | 0 (0.0%) | 0 (0.0%) |
| Other / Unknown | 4 (12.1%) | 2 (1.4%) | 0 (0.0%) | 2 (3.3%) | 0 (0.0%) | 0 (0.0%) |

### Clustering with molecular features

iPSC-derived spinal motor neuron transcriptomic data from day 60 were analyzed using five dimensionality-reduction methods followed by K-means clustering to identify molecularly distinct ALS subgroups. Clustering performance was evaluated using the Davies–Bouldin and Calinski–Harabasz indices, and the optimal cluster number was selected by the elbow method.

Molecular data gathered at day 60 of the iPSN^12^ was selected for downstream data analyses because it showed both the greatest number of differentially expressed mRNA species (out of the 18 TDP-43 loss of function related targets) and the largest effect sizes, as reflected by the most significant p-values relative to control subjects6^11^. Additionally, data was not stratified by sex after finding no significant differences in the genetic expression of male and female patients at that time point. Of the five dimensionality reduction methods, spectral embedding paired with K-means clustering displayed the lowest DB index of 0.34 and the highest CH index of 623 (Table 2). Based on the elbow plot of explained variance (Fig. 1a) as well as additional examination of alternative cluster numbers, four clusters was selected as the ideal number of subgroups. With this labeling, all 12 patients with the C9orf72 mutation naturally grouped into a single cluster, thereafter, known as Subgroup C. Meanwhile, the sALS group split into three clusters, which were labeled as the S1 (n=51), S2 (n=60), and S3 (n=18) subgroups (Fig. 1b). Adding in control patients while clustering with k=4 resulted in the same subgroups, but with the S3 and control patients grouped together, indicating similar molecular expression.

**Table 2.**
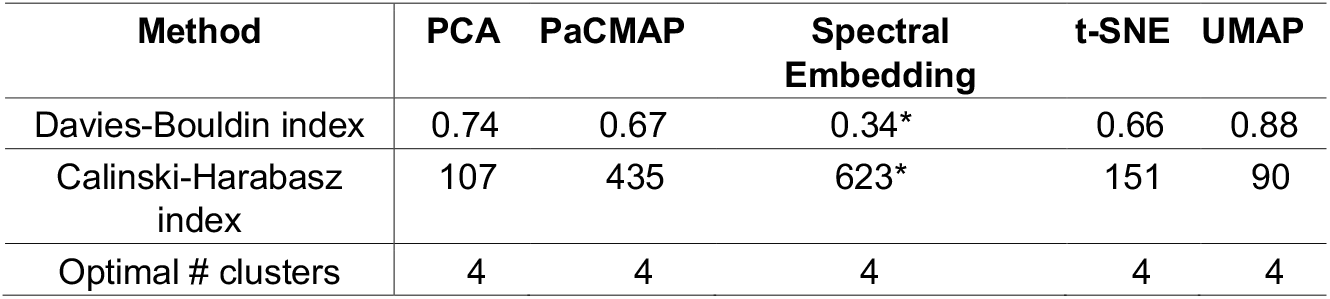
Comparison of dimensionality reduction methods for clustering of molecular features. Asterisked values represent the best performance metrics: the lowest Davies-Bouldin index and highest Calinski-Harabasz index.

**Figure 1.**
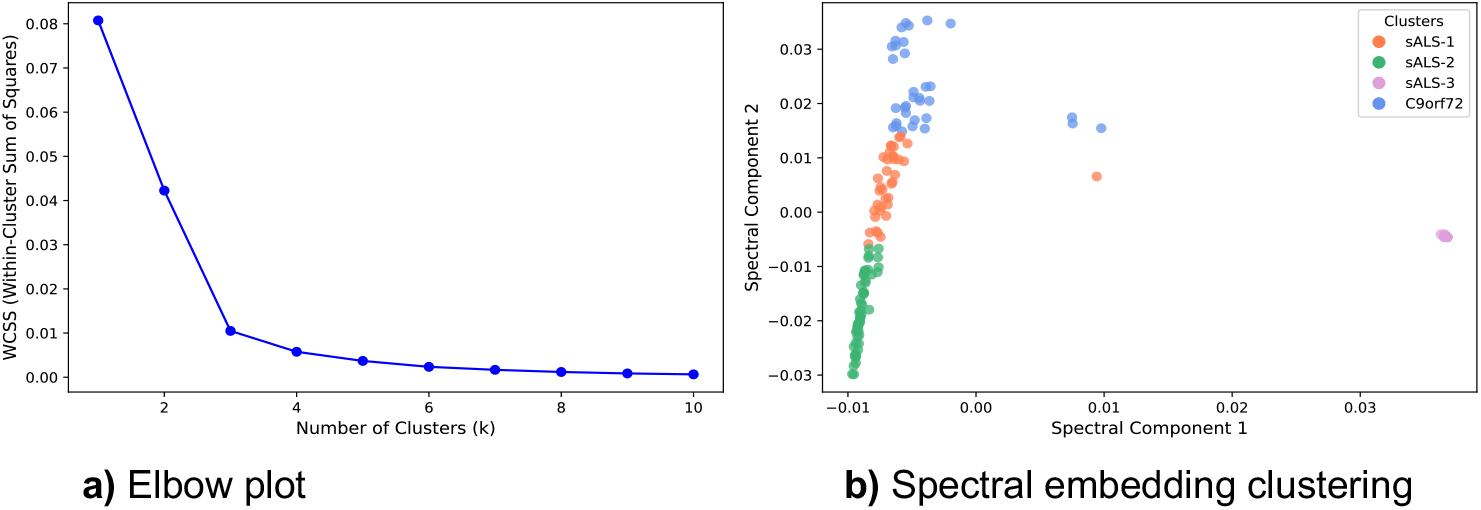
Data-driven clustering pipeline for ALS patient stratification. (a) Elbow plot showing WCSS across increasing numbers of clusters, with an inflection point supporting selection of four clusters. (b) Visualization of the final clusters in reduced-dimensional space with spectral embedding and K-means.

**Figure 2.**
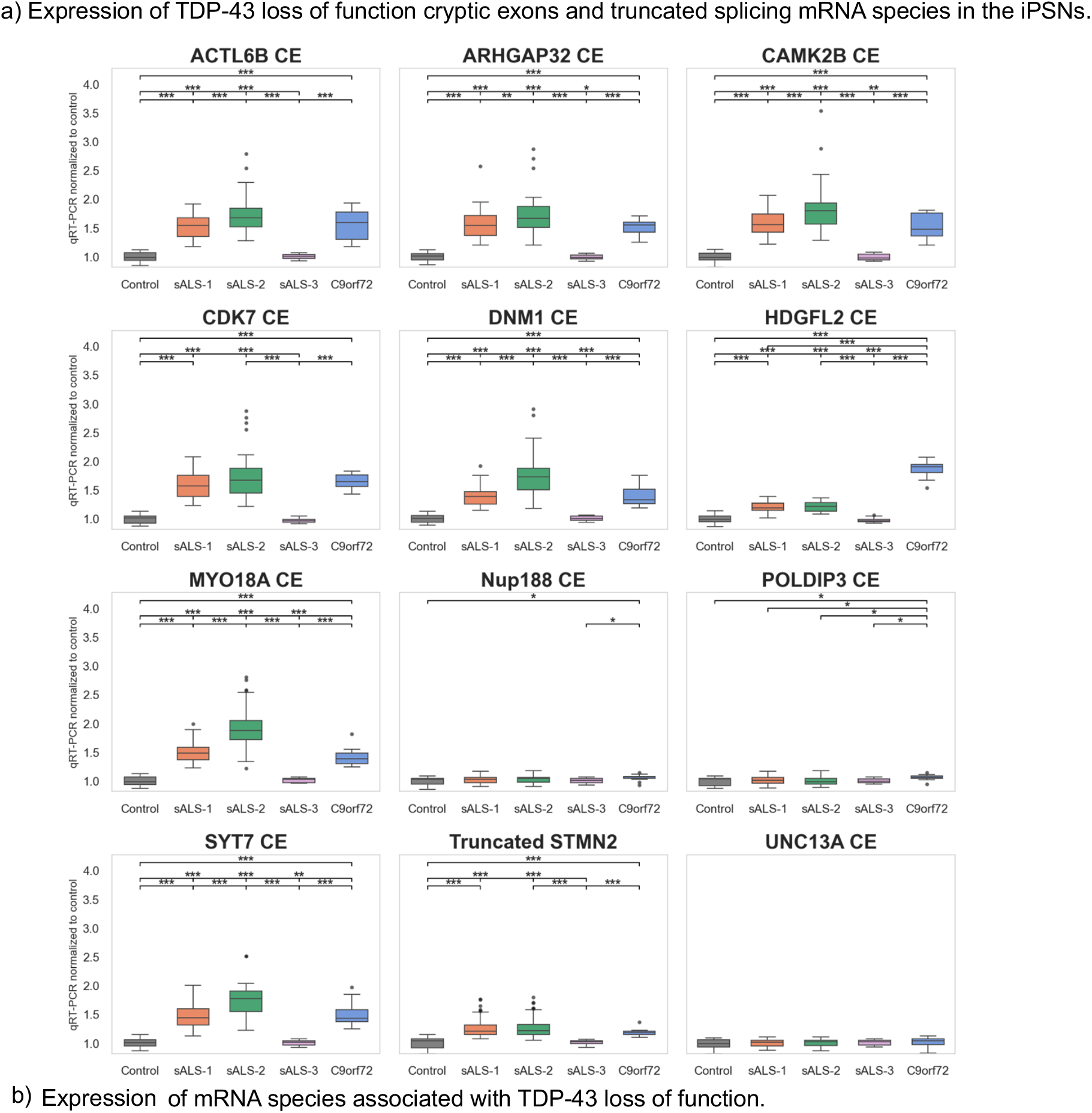
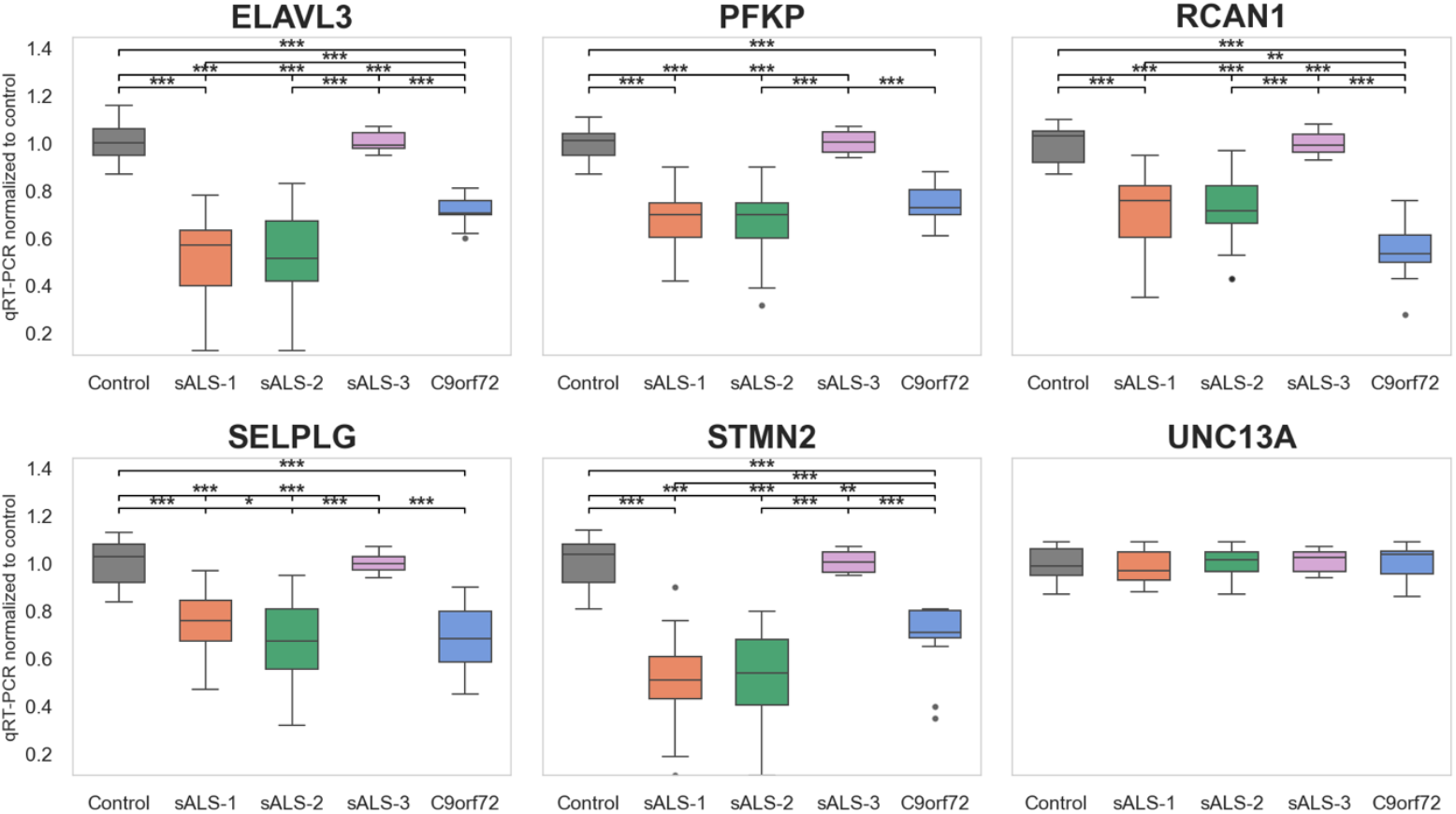
**A)** Expression of TDP-43 loss of function cryptic exons and truncated splicing mRNA species in the iPSNs. **B)** Expression of mRNA species associated with TDP-43 loss of function. Pairwise cluster differences were assessed using Wilcoxon rank-sum tests (* p < 0.05, ** p < 0.01, *** p < 0.001). Significance bars are grouped by comparison width: comparisons between adjacent clusters appear on the bottom tier, while comparisons spanning more distant clusters appear on higher tiers.

### Molecular Differences between ALS subgroups

Expression and splicing values for 18 TDP-43–related transcripts were compared across subgroups using Kruskal–Wallis tests, followed by pairwise Wilcoxon Rank-Sum tests for significant features. These analyses evaluated subgroup-level variation in TDP-43 loss-of-function signatures.

Seventeen genes were found with significantly different expression profiles (p < 0.05) across patients in the S1, S2, S3, C, and control groups (Fig. 2). TDP-43 CE expression was consistently lowest in non-ALS control patients and significantly elevated in one or more ALS subgroups (Fig. 2a). Specifically, MYO18A TDP-43 CE (p=3.92e-28), DNM1 CE (p=2.30e-26), HDGFL2 CE (p=6.69e-25), SYT7 CE (p=2.61e-25), and CAMK2B CE (p=1.46e-24) showed increased CE inclusion. Other genes with elevated CE levels included ACTL6B (p=1.23e-23), ARHGAP32 (p=3.28e-23), CDK7 (p=1.07e-22), Truncated STMN2 (5.68e-21), POLDIP3 (p=4.04e-2), and NUP188 (p=3.38e-2). Simultaneously, known TDP-43 target RNAs, comprising ELAVL3 (p=3.22e-24), RCAN1 (p=3.70e-22), STMN2 (1.49e-23), SELPLG (5.14e-21), PFKP (p=9.87e-23), and UNC13A (p=4.94e-2), were downregulated in ALS patients relative to controls (Fig. 2b). These expression changes were most pronounced in the S2 subgroup, which showed the greatest increases in TDP-43 CE expression and strongest decreases in TDP-43 target transcripts.

One-vs-rest subgroup discrimination based on TDP-43–associated cryptic exon (CE) expression was next assessed (Table 3). For S2 patients, subgroup membership could be identified with near-perfect accuracy using a combination of four CE events: MYO18A CE, DNM1 CE, SYT7 CE, and ACTL6B CE (AUC = 0.993). For subgroups S3 and C, a single gene expression or mRNA splicing event was sufficient to achieve perfect separation from the remaining samples (AUC = 1.0). These events were STMN2 for S3 and HDGFL2 CE for group C. In contrast, S1 showed lower discrimination performance (AUC ≈ 0.67). Although STMN2 was the most informative feature for this subgroup, no small subset of CE events achieved robust discrimination.

**Table 3.** Discriminative cryptic exon features across ALS molecular subgroups. Summary of subgroup discrimination performance based on TDP-43–associated CE expression, including one-vs-rest classification performance, the number of CE features selected by sparse classification, the most strongly discriminative CE events, and the minimal CE subset required to achieve peak discrimination. Classification performance is reported as cross-validated area under the receiver operating characteristic curve (AUC).

| Subgroup | One-vs-rest AUC | # CEs selected | Minimal CEs for peak AUC | Top CEs | Peak AUC |
| --- | --- | --- | --- | --- | --- |
| sALS-1 | 0.773 | 10 | 1 | STMN2 | 0.669 |
| sALS-2 | 0.978 | 8 | 4 | MYO18A_CE, DNM1_CE, SYT7_CE, ACTL6B_CE | 0.993 |
| sALS-3 | 1.000 | 10 | 1 | STMN2 | 1.000 |
| C9orf72 | 1.000 | 1 | 1 | HDGFL2_CE | 1.000 |

### Clinical correlates

Longitudinal clinical outcomes were analyzed with LMMs including time, subgroup, and their interaction as fixed effects and patient ID as a random effect. Significant time–cluster interaction terms indicated subgroup-specific progression rates.

95 longitudinal clinical outcomes were examined, spanning the categories of: Revised ALS Functional Rating Scale (ALSFRS-R), deep tendon reflex scores, HHD measurements, cognitive-behavioral screening, Ashworth spasticity scores, grip strength, auxiliary bloodwork markers, and vital capacity volumes.

36 clinical measures were found to have significantly different rates of progression between clusters after LMM analysis. The average rates of progression over one month (four weeks) are reported in Table 4, with significant pairwise differences in Table 5.

**Table 4.**
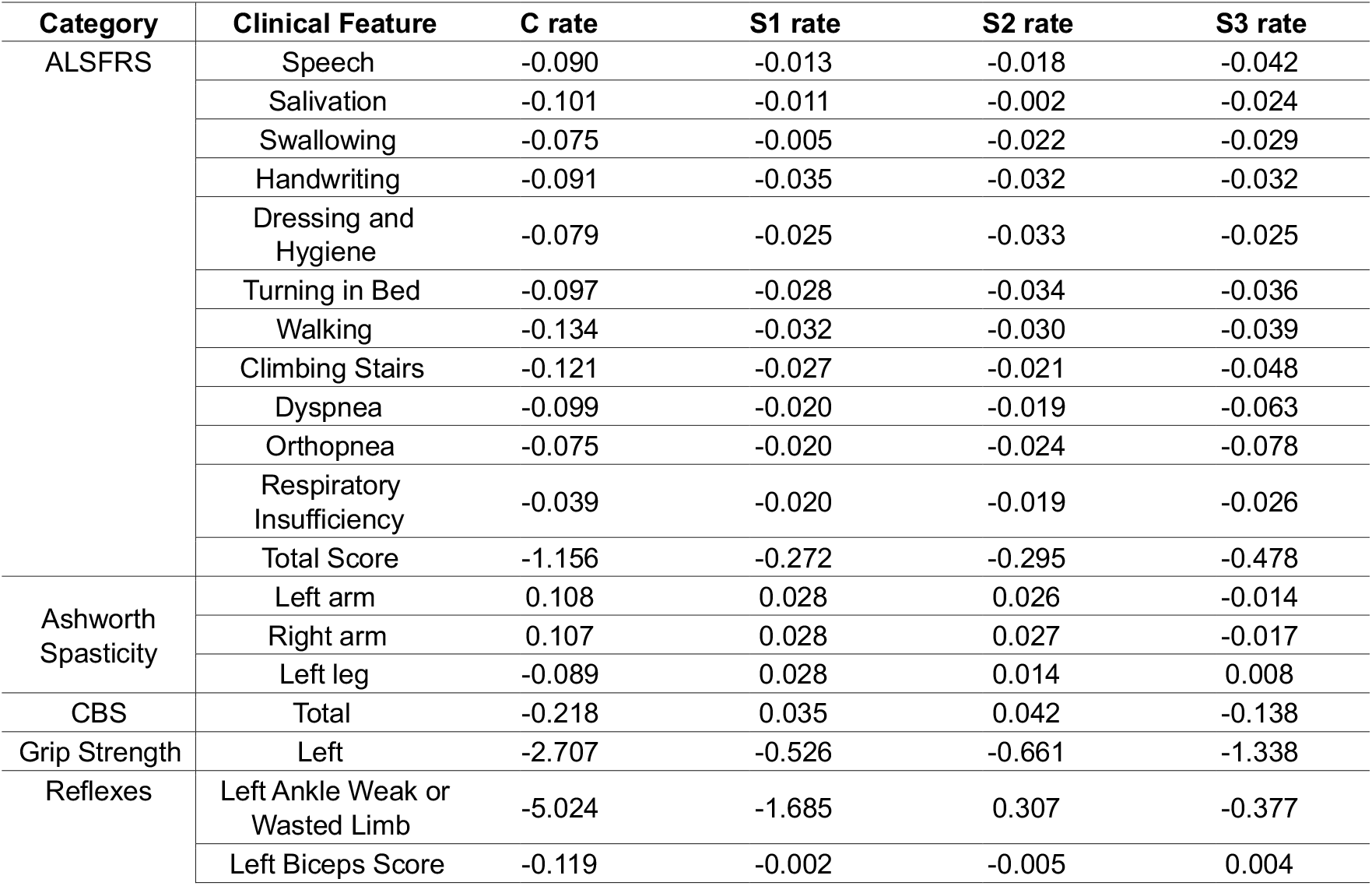
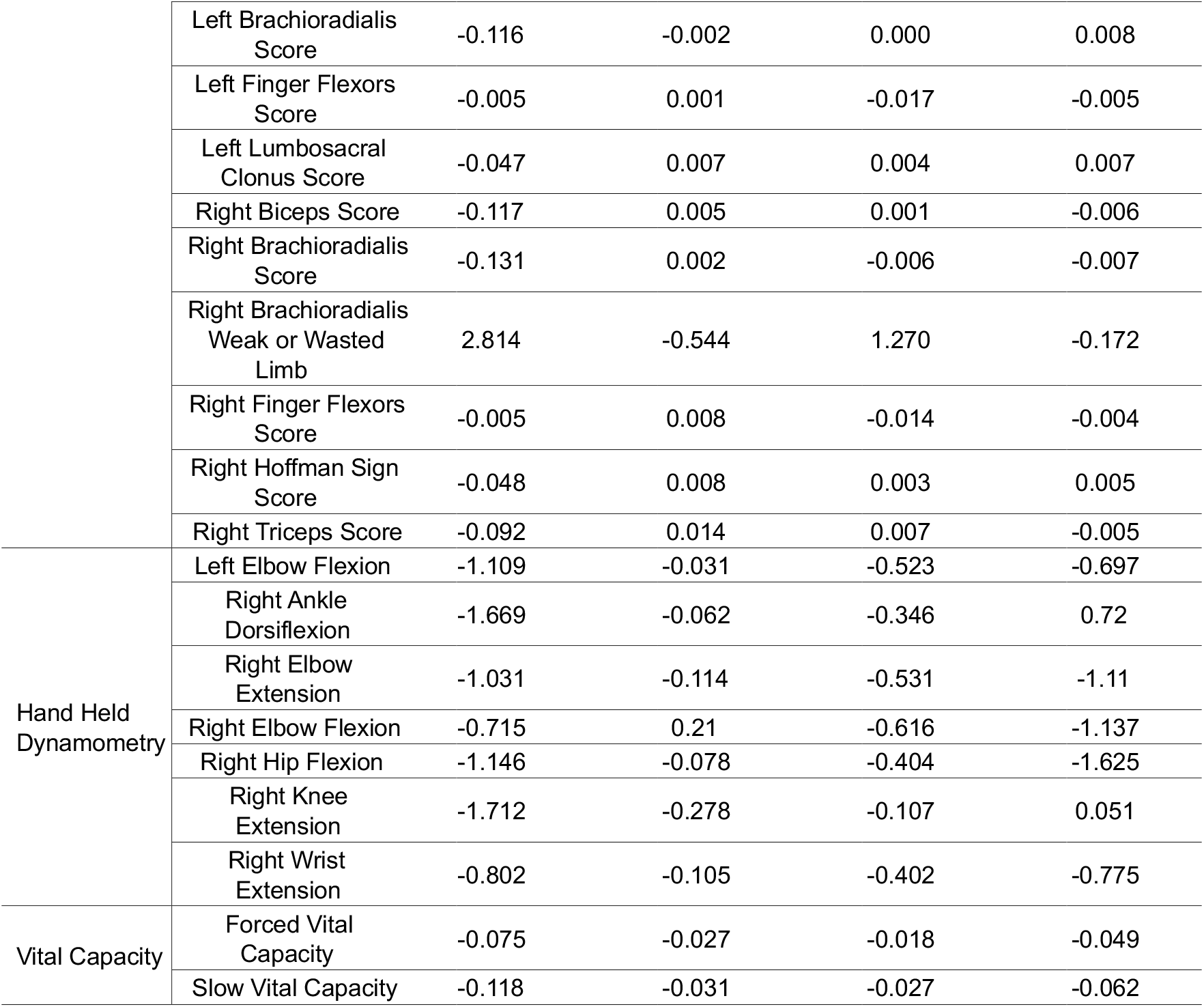
Rate of change in clinical features over 4 weeks (28 days), estimated from LMMs. Values reflect interaction effects between time and cluster, representing subgroup-specific progression rates.

| Category | Clinical Feature | C rate | S1 rate | S2 rate | S3 rate |
| --- | --- | --- | --- | --- | --- |
| ALSFRS | Speech | -0.090 | -0.013 | -0.018 | -0.042 |
|  | Salivation | -0.101 | -0.011 | -0.002 | -0.024 |
|  | Swallowing | -0.075 | -0.005 | -0.022 | -0.029 |
|  | Handwriting | -0.091 | -0.035 | -0.032 | -0.032 |
|  | Dressing and Hygiene | -0.079 | -0.025 | -0.033 | -0.025 |
|  | Turning in Bed | -0.097 | -0.028 | -0.034 | -0.036 |
|  | Walking | -0.134 | -0.032 | -0.030 | -0.039 |
|  | Climbing Stairs | -0.121 | -0.027 | -0.021 | -0.048 |
|  | Dyspnea | -0.099 | -0.020 | -0.019 | -0.063 |
|  | Orthopnea | -0.075 | -0.020 | -0.024 | -0.078 |
|  | Respiratory Insufficiency | -0.039 | -0.020 | -0.019 | -0.026 |
|  | Total Score | -1.156 | -0.272 | -0.295 | -0.478 |
| Ashworth Spasticity | Left arm | 0.108 | 0.028 | 0.026 | -0.014 |
|  | Right arm | 0.107 | 0.028 | 0.027 | -0.017 |
|  | Left leg | -0.089 | 0.028 | 0.014 | 0.008 |
| CBS | Total | -0.218 | 0.035 | 0.042 | -0.138 |
| Grip Strength | Left | -2.707 | -0.526 | -0.661 | -1.338 |
| Reflexes | Left Ankle Weak or Wasted Limb | -5.024 | -1.685 | 0.307 | -0.377 |
|  | Left Biceps Score | -0.119 | -0.002 | -0.005 | 0.004 |
|  | Left Brachioradialis Score | -0.116 | -0.002 | 0.000 | 0.008 |
|  | Left Finger Flexors Score | -0.005 | 0.001 | -0.017 | -0.005 |
|  | Left Lumbosacral Clonus Score | -0.047 | 0.007 | 0.004 | 0.007 |
|  | Right Biceps Score | -0.117 | 0.005 | 0.001 | -0.006 |
|  | Right Brachioradialis Score | -0.131 | 0.002 | -0.006 | -0.007 |
|  | Right Brachioradialis Weak or Wasted Limb | 2.814 | -0.544 | 1.270 | -0.172 |
|  | Right Finger Flexors Score | -0.005 | 0.008 | -0.014 | -0.004 |
|  | Right Hoffman Sign Score | -0.048 | 0.008 | 0.003 | 0.005 |
|  | Right Triceps Score | -0.092 | 0.014 | 0.007 | -0.005 |
| Hand Held Dynamometry | Left Elbow Flexion | -1.109 | -0.031 | -0.523 | -0.697 |
|  | Right Ankle Dorsiflexion | -1.669 | -0.062 | -0.346 | 0.72 |
|  | Right Elbow Extension | -1.031 | -0.114 | -0.531 | -1.11 |
|  | Right Elbow Flexion | -0.715 | 0.21 | -0.616 | -1.137 |
|  | Right Hip Flexion | -1.146 | -0.078 | -0.404 | -1.625 |
|  | Right Knee Extension | -1.712 | -0.278 | -0.107 | 0.051 |
|  | Right Wrist Extension | -0.802 | -0.105 | -0.402 | -0.775 |
| Vital Capacity | Forced Vital Capacity | -0.075 | -0.027 | -0.018 | -0.049 |
|  | Slow Vital Capacity | -0.118 | -0.031 | -0.027 | -0.062 |

**Table 5.** Significant clinical features based on LMM analyses. Pairwise comparisons were obtained by rotating the reference level to each cluster in turn. Values represent estimated differences in mean rates of progression per month (4 weeks) between the two groups, with p-values in parentheses. Comparisons that did not reach statistical significance (p-value > 0.05) are denoted by “-”.

| Category | Clinical Feature | C vs. S1 | C vs. S2 | C vs. S3 | S1 vs. S2 | S1 vs. S3 | S2 vs. S3 |
| --- | --- | --- | --- | --- | --- | --- | --- |
| ALSFRS | Speech | 0.077<br>(<0.001) | 0.071<br>(<0.001) | 0.048<br>(0.020) | -0.029<br>(0.002) | -0.024<br>(0.012) | — |
|  | Salivation | 0.089<br>(<0.001) | 0.099<br>(<0.001) | 0.077<br>(0.001) | — | — | — |
|  | Swallowing | 0.070<br>(0.001) | 0.054<br>(0.010) | 0.047<br>(0.042) | -0.016<br>(0.002) | -0.023<br>(0.031) | — |
|  | Handwriting | 0.057<br>(0.027) | 0.060<br>(0.019) | 0.060<br>(0.032) | — | — | — |
|  | Dressing and Hygiene | 0.054<br>(0.027) | — | 0.054<br>(0.042) | — | — | — |
|  | Turning in bed | 0.069<br>(0.007) | 0.062<br>(0.013) | 0.061<br>(0.028) | — | — | — |
|  | Walking | 0.103<br>(<0.001) | 0.105<br>(<0.001) | 0.095<br>(<0.001) | — | — | — |
|  | Climbing Stairs | 0.094<br>(0.001) | 0.100<br>(0.001) | 0.073<br>(0.022) | — | — | — |
|  | Dyspnea | 0.079<br>(0.007) | 0.079<br>(0.006) | — | — | -0.043<br>(0.003) | -0.044<br>(0.002) |
|  | Orthopnea | — | — | — | — | -0.058<br>(0.001) | -0.054<br>(0.001) |
| | Total score | 0.883<br>($<0.001$) | 0.860<br>($<0.001$) | 0.678<br>($<0.001$) | -0.206<br>(0.010) | -0.183<br>(0.019) | — |
| Ashworth | Left arm | — | — | -0.122<br>(0.025) | — | — | — |
|  | Right arm | — | — | -0.124<br>(0.029) | — | — | — |
|  | Left leg | 0.117<br>(0.027) | — | — | — | — | — |
| CBS | Total | 0.253<br>(0.008) | 0.260<br>(0.006) | — | — | -0.173<br>($<0.001$) | -0.180<br>($<0.001$) |
| Grip Strength | Left | 2.180<br>(0.016) | 2.046<br>(0.022) | — | — | — | — |
| HHD | Left Elbow Flexion | 1.078<br>(0.040) | — | — | — | — | — |
|  | Right Ankle Dorsiflexion | 1.607<br>(0.017) | 1.323<br>(0.044) | 2.389<br>(0.001) | — | — | 1.066<br>(0.014) |
| | Right Elbow Extension | 0.917<br>(0.020) | — | — | -0.417<br>(0.023) | -0.996<br>($<0.001$) | -0.580<br>(0.034) |
| | Right Elbow Flexion | — | — | — | -0.826<br>(0.001) | -1.346<br>($<0.001$) | — |
|  | Right Hip Flexion | — | — | — | -1.547<br>(0.001) | — | -1.220<br>(0.002) |
|  | Right Knee Extension | 1.434<br>(0.025) | 1.605<br>(0.011) | 1.763<br>(0.012) | — | — | — |
|  | Right Wrist Extension | — | — | — | -0.291<br>(0.038) | -0.664<br>(0.008) | — |
| Reflexes | Left Ankle Weak/Wasted | 4.647<br>(0.047) | — | 5.331<br>(0.026) | — | — | 1.991<br>(0.040) |
|  | Left Biceps | 0.117<br>(0.005) | 0.114<br>(0.006) | 0.124<br>(0.004) | — | — | — |
|  | Left Brachioradialis | 0.114<br>(0.018) | 0.116<br>(0.017) | 0.123<br>(0.014) | — | — | — |
|  | Left Finger Flexors | — | — | — | -0.0.18<br>(0.042) | — | — |
|  | Left Lumbosacral Clonus | -0.037<br>(0.076) | — | — | 0.015<br>(0.094) | 0.021<br>(0.031) | — |
|  | Right Biceps | — | 0.118<br>(0.005) | 0.110<br>(0.010) | — | — | — |
|  | Right Brachioradialis | 0.133<br>(0.007) | 0.124<br>(0.012) | 0.012<br>(0.016) | — | — | — |
|  | Right Brachioradialis Weak/Wasted | — | — | — | 1.814<br>(0.042) | — | — |
|  | Right Finger Flexors | — | — | — | -0.021<br>(0.020) | — | — |
|  | Right Hoffman Sign | 0.056<br>(0.004) | 0.051<br>(0.010) | 0.053<br>(0.010) | — | — | — |
|  | Right Triceps | 0.106<br>(0.014) | 0.099<br>(0.022) | 0.087<br>(0.048) | — | — | — |
| Vital Capacity | Forced Vital Capacity | — | — | — | — | — | -0.030<br>(0.014) |
| | Slow Vital Capacity | 0.087<br>($<0.001$) | 0.090<br>($<0.001$) | 0.056<br>(0.030) | — | — | -0.034<br>(0.044) |

#### ALSFRS

ALSFRS-R or simply ALSFRS is a validated clinical measure of ALS progression, with lower scores indicating greater functional impairment^28^. The total score, an ordinal scale-based scoring, comprised of 12 subscores evaluating bulbar, motor, and respiratory function, provides a summary of patient disability. LMMs revealed subgroup-specific rates of decline over 4 weeks, with subgroup C showing the steepest drop in ALSFRS total score (p<0.001), followed by S3, S2, and S1 (Table 4). Pairwise comparisons of the interaction effects (i.e., rate differences between subgroups) identified significant differences across multiple ALSFRS subscores (Table 5). In particular, speech, swallowing, walking, and climbing stairs showed significantly faster decline in subgroup C compared to others. S3 showed greater decline compared to S1 and S2, with relatively larger drops in orthopnea (−0.078 points/month), dyspnea (−0.063 points/month), and speech (−0.042 points/month).

#### Ashworth

Ashworth spasticity scores measure a patient’s increase in muscle tone since the previous visit, with 0 indicating no increase and 4 indicating significant increase^29^. Greatly increased muscle tone to the point of spasticity points to an inability to stop muscle contraction and is a unique marker of upper motor neuron dysfunction. Three of four Ashworth spasticity scores were found significant: left arm, right arm, and left leg (Table 4). The difference between rates of progression was significant between subgroups C and S3 in left (p=0.025) and right arm (p=0.029) measures, with subgroup C having the greatest rate of increase, and S3 showing decreased scores (Table 4). In the left leg, spasticity scores were significantly lower in C than S1 (p=0.027). No other pairwise differences reached statistical significance.

#### Cognitive Behavioral Screen

The ALS Cognitive Behavioral Screen (CBS) measures cognitive and behavioral ability (concentration, retrieval, attention, etc.) in ALS patients, with lower scores indicating cognitive and behavioral impairment^30^. Clusters C and S3 showed overall decreases in CBS scores, with rates of progression significantly lower than those of S1 and S2.

#### Grip Strength

Grip strength testing measures force exerted by one’s hands and forearms in pounds^31^. Significant declines in left grip strength were observed for the reference cluster C, with an average decrease of −2.71 lbs/month (Table 3). Subgroups S1 and S2 had significantly slower rates of decline than group C, averaging −0.53 (p=0.016) and −0.66 (p=0.022) lbs/month, respectively (Table 4 and Figure 3c).

**Figure 3.**
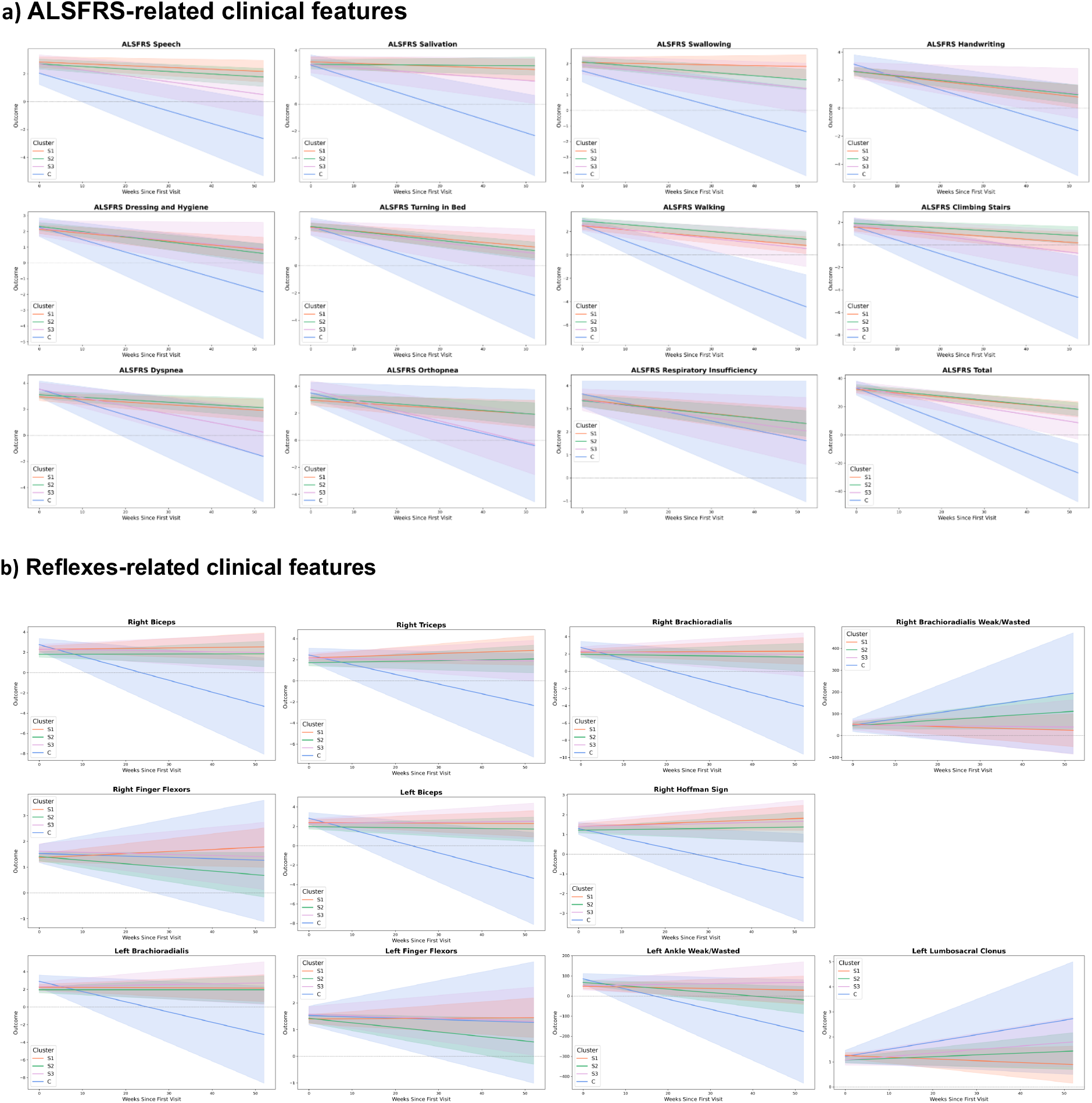
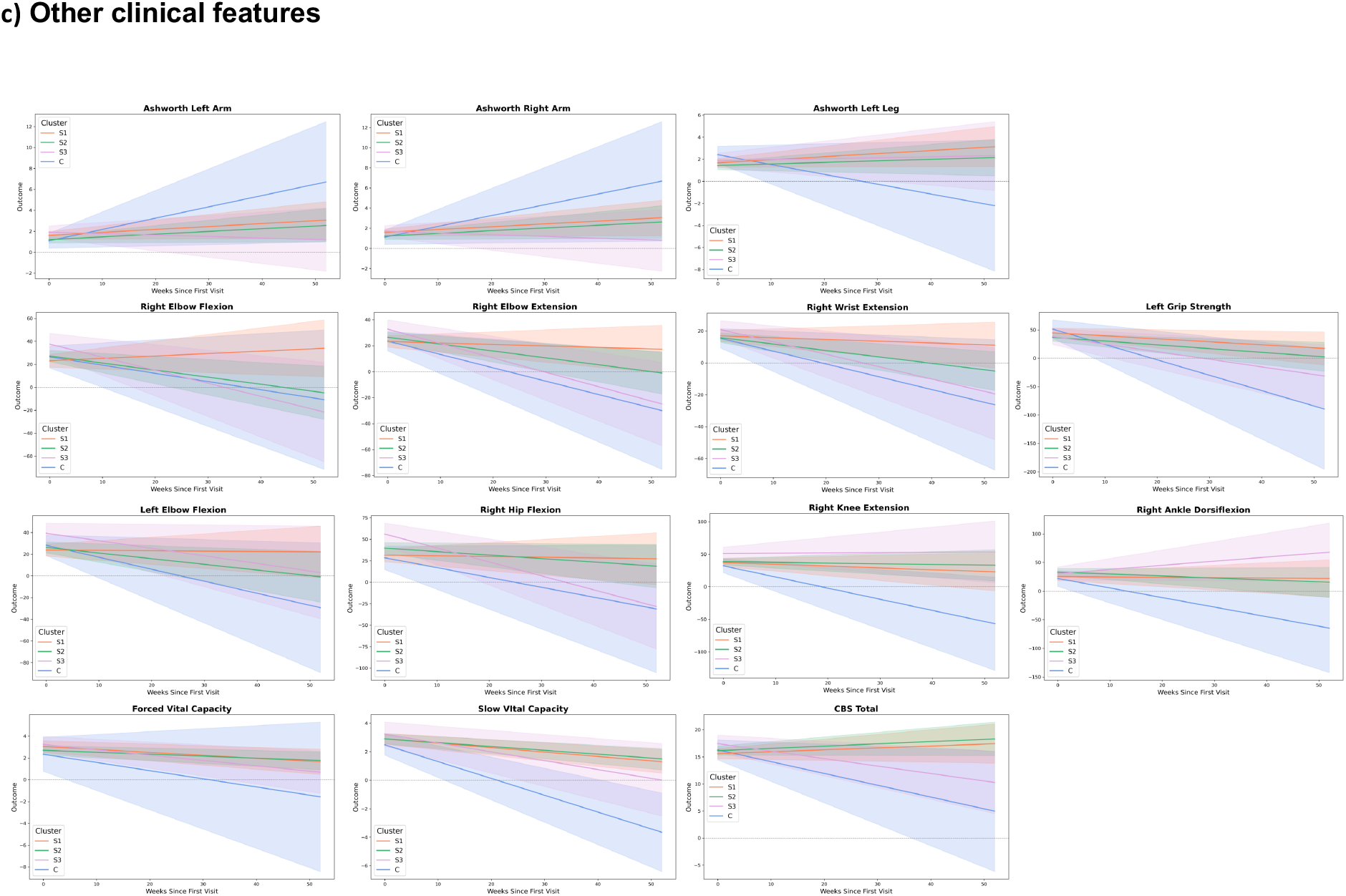
Longitudinal trajectories of clinical features across clusters, grouped by clinical domains including a) ALSFRS, b) deep tendon reflexes and c) other clinical features (e.g. strength, forced vital capacity, cognition).

#### Hand-held Dynamometry

HHD scores are used to measure muscle strength, with scores reflecting the amount of generated force in pounds^24^. Seven categories were found to have significant differences in progression among subgroups. LMM slope values revealed that cluster C generally had the fastest rate of decline, leading in five of seven measures. In contrast, S1 is generally the slowest to decline and even improves in right elbow flexion.

Between-cluster interaction effects mirror these patterns. The largest number of significant differences occur for C vs. S1, with S1 having a significantly slower trajectory especially in right ankle dorsiflexion (p=0.017) and right elbow extension (p=0.02) (Table 5). The same pattern was observed with S1 and S2, with S1 having a significantly slower trajectory in scores for right elbow extension (p=0.023), right elbow flexion (p=0.001), and right hip flexion (p=0.001).

#### Reflexes

Reflex scores assess the integrity of the central and peripheral nervous systems by quantifying involuntary muscle responses. Abnormal reflexes, including hyperreflexia or absence of reflexes, can indicate upper or lower motor neuron dysfunction in ALS^32^. Eight reflex-related measures were found to be significant: left ankle (weak or wasted limb), left biceps, left brachioradialis, right biceps, right brachioradialis, right triceps, and Hoffman sign scores bilaterally (Table 4).

Based on LMM interaction effects, cluster C showed a consistent decline in reflex scores across all eight measures, with the largest drop observed in left ankle weakness and the smallest changes in Hoffman sign scores. In contrast, subgroups S1, S2, and S3 generally showed no significant changes over time for most reflexes. Cluster C showed significantly lower scores than S1, S2, and S3 for left biceps, left brachioradialis, right brachioradialis, right Hoffman sign, and right triceps.

Left ankle weakness showed a notable decline in both cluster C and S1, whereas S3 exhibited a slight increase. For right triceps, reflex scores slightly increased over time in S1 and S2, with negligible changes in S3 and C.

Group-wise comparisons of model-estimated coefficients are shown in Figure 3b and detailed in Table 4. Among all reflex measures, biceps scores on both sides were the most consistently significant (p < 0.01 across all S groups), while right brachioradialis weakness was uniquely elevated in C.

#### Vital Capacity

Slow vital capacity (SVC) is the volume of air, in liters, that a patient can exhale slowly from total lung capacity to residual volume. Forced vital capacity (FVC) is the maximal volume of air, in liters, that can be exhaled forcefully after a full inspiration. Vital capacity is a standard pulmonary function measure in ALS and is used to track respiratory decline over time^33^. Both SVC and FVC differed significantly across patient subgroups (Table 4). Cluster C showed the steepest rate of SVC decline, progressing significantly faster than the other three clusters, and S3 declined faster than S2 in both SVC and FVC.

## Discussion

The clinical and molecular heterogeneity of ALS has driven extensive efforts to identify patient subgroups based on phenotype, clinical prognosis, and molecular data^4,7,34–38^. Accurate stratification of patients could provide insight into the underlying pathological mechanisms of the disease, allowing for focused therapeutic innovation as well as improved clinical trial design. Ultimately, targeted therapies for specific ALS hallmarks could be developed for personalized treatment strategies depending on ALS subtype.

In this study, data-driven machine learning methods were leveraged to identify molecular ALS subtypes based on transcriptomic markers of TDP-43 dysfunction. To date, it is known that the vast majority of sporadic ALS patient have TDP-43 pathology, typified as cytoplasmic and sometimes, nuclear aggregates. But the functional loss of TDP-43, as reflected by measure of altered mRNA and cryptic RNA species, has heretofore not been quantified across multiple molecular species and multiple patients. The large panel of Answer ALS iPS cell lines, with matched molecular and clinical data, provide the first large scale population-based approach to this analytic. Since the IPS model system appears to closely match actual human brain, this model system provides an opportunity to evaluate if the patterns of TDP-43 loss of function can elucidate different molecular subgroups of actual patients. To that end, spectral embedding and K-means clustering were applied to qRT-PCR data from 141 sALS patient iPSNs, including those with and without C9orf72 mutations. With optimal clustering creating four clusters, patients with the C9orf72 mutation were effectively isolated from other sALS patients. The remaining population was further divided into three groups (S1, S2, and S3). Interestingly, the S3 patients clustered closely with non-ALS control patients, suggesting minimal TDP-43 dysfunction at the molecular level. Consistent with this clustering structure, subgroup discrimination based on the qRT-PCR data revealed heterogeneity in TDP-43 dysfunction across sALS subtypes. While subgroups S3 and C exhibited highly specific CE signatures--each defined by just one dominant CE event—S2 required a coordinated set of four CE changes for accurate identification, and S1 showed only modest discriminability. Notably, the gene expression change in STMN2 was the top discriminating feature for both S1 and S3, potentially highlighting an area for future research.

LMMs were then used to analyze cluster-wise differences in longitudinal clinical outcomes. Of 95 potential clinical correlates, 36 measures in the categories of ALSFRS subscores, muscle strength (via HHD and grip strength), reflex changes, Ashworth spasticity ratings, and vital capacity were found to be significantly different between subgroups.

Subgroups S1 and S2 represented the mild end of the clinical spectrum, with slower functional decline and largely preserved limb strength. Both demonstrated modest declines in CBS scores, whereas S2 showed subtle indicators of lower motor neuron involvement. These findings suggest that S1 and S2 may reflect subtypes with relatively restricted or slowly evolving pathology.

S3 displayed a distinctly heterogeneous clinical pattern, reflecting uneven vulnerability across motor and respiratory systems. Larger declines in orthopnea and speech point to more pronounced involvement of bulbar and respiratory pathways, while slight reductions in arm tone and mixed HHD trajectories suggest that compensatory mechanisms may persist in some regions despite accelerated degeneration in others. S3 also exhibited greater respiratory decline than S2, placing it clinically between the more stable S1/S2 groups and the rapidly progressive C group. Declines in CBS scores further support the interpretation that S3 represents a subtype with broader multisystem impairment.

Cluster C represents the most aggressive clinical phenotype, consistent with its marked molecular divergence. This observation aligns with prior studies showing that C9orf72 repeat expansion is associated with a more severe disease course and shorter survival^39, 40^. Notably, our identification of this subgroup emerged purely from transcriptomic clustering, suggesting that C9orf72 status exerts a sufficiently strong molecular signature to be recovered in an unsupervised framework.

Patients in this subgroup experienced steep declines across ALSFRS mobility, bulbar, and respiratory functions, suggestive of widespread and rapidly advancing neurodegeneration. Pronounced increases in Ashworth spasticity in both arms emphasize substantial upper motor neuron involvement, while decreased left-leg tone, opposite the trend seen in other subgroups, highlights region-specific differences in disease expression. C also exhibited the largest reductions in grip strength and HHD measures, along with clear reflex deterioration and increases in weak/wasted items, reflecting simultaneous loss of upper and lower motor neuron integrity. The steepest declines in SVC and FVC further underscore the severe respiratory vulnerability of this subgroup, distinguishing C as the most clinically and physiologically compromised cluster.

These clinical findings reveal that molecular subtyping based on TDP-43 dysfunction aligns with distinct patterns of functional, tone, strength, and reflex change. Cluster C appears to comprise of patients with a rapid and globally progressing disease with broad strength loss. S1 seems to be more slowly progressive, and S2 and S3 as intermediate with mixed trajectories of muscle strength and functionality. In particular, the S3 profile may indicate alternative mechanisms (e.g., non-canonical pathways) that are not fully captured by the TDP-43 markers from this study.

For clinical trial design, the presence of S3 patients would have critical implications. Such individuals may be less responsive to therapies targeting TDP-43, and their inclusion could confound trial outcomes. However, for other sALS patients with marked TDP-43 dysfunction, further research should be conducted to assess how their increased levels of cryptic exons inclusion in target transcripts and decreased levels of known TDP-43 target RNAs are related to clinical symptomatology.

While this study focuses on grouping by TDP-43 loss of function, the data is derived from iPSN models, which may not fully capture the multidimensionality of ALS in vivo. Although recent studies note the close similarity between iPS TDP-43 and autopsy tissue loss of function determination, especially those rare examples with matched IPS lines and postmortem tissue, this close correlation of TDP-43 model readout and same patient tissue analytics warrant further future investigation with larger set of matched iPS cell lines and postmortem tissue. Additionally, variability in clinical data, especially in terms of subgroup representation, may limit generalizability. Some subgroups had relatively small sample sizes, potentially caused by earlier mortality, leading to possible underrepresentation of more aggressive disease presentations.

Future work should focus on the relationships between the molecular profiles of the subgroups and their outward clinical presentations. More research is needed to identify specific molecular pathways contributing to physical symptoms. Additionally, patients falling in the S3 cluster, with minimal TDP-43 dysfunction but substantial declines in strength, should be studied for alternative and additional ALS mechanisms. Finally, we hope to validate our clusters in future studies by prospectively labeling patients with their clusters and predicting clinical outcomes.

## Conclusions

This study developed an ML-based framework that used transcriptomic and splicing signatures from iPSC-derived motor neurons to identify biologically meaningful subgroups of sALS. We defined four molecularly distinct clusters: S1, S2, S3, and C, which exhibited clear differences in disease progression. S3 showed rapid global decline but unexpectedly preserved motor strength early on, suggesting possible compensatory mechanisms; S1 demonstrated localized left ankle weakness with mixed progression; and S2 showed reduced reflex and spasticity scores consistent with predominant lower motor neuron involvement. These findings highlight how molecular subgrouping can help clarify ALS heterogeneity, refine prognosis, and guide more targeted therapeutic development and clinical-trial design. Continued validation of these subgroups and deeper investigation of their underlying pathways will be critical for advancing precision medicine in ALS.

## Declarations

## Ethics approval and consent to participate

All iPSC lines used within this study were deidentified and collected under standard IRB approved consents from each of the participating sources. The Johns Hopkins IRB protocols that are represented in this study include: Answer ALS: Individualized Initiative for ALS Discovery, IRB00082277; Answer Amyotrophic Lateral Sclerosis (AALS) Research Data Portal, IRB00392828.

## Consent for publication

Not applicable.

## Availability of data and materials

The datasets analyzed during the current study are available at https://dataportal.answerals.org/home. Additional raw data will be shared by the corresponding author upon request.

## Competing interests

The authors declare no competing interests relevant to the observations in this study.

## Funding

This work was supported by The Robert Packard Center for ALS Research at Johns Hopkins, and NIH (R35NS132179; R01NS122236).

## Author contributions statement

TC, ST, YG, PV, RL, and MP performed computational analyses, interpreted results, and wrote the manuscript. JR and AC conceived the experiment, provided data, and reviewed the proposed protocol. JG, CT, AY, NS, CH, and IV reviewed the technical protocol and provided feedback. All authors reviewed and approved the manuscript.

## Acknowledgments

The authors wish to acknowledge Brian Scott Caffo and Jiangxia Wang for their guidance and support in planning the data analysis workflow. The overall program would not have been posible without the Answer ALS Research Program and its comprehensive collection of sALS and fALS patient derived iPS cell lines. We also sincerely thank the ALS patients and their families whose contributions made this work possible.

## Notes

### Competing Interest Statement

The authors have declared no competing interest.

### Summary of Updates

Subgroup discrimination analyses were now performed to further assess whether differences in specific TDP-43 associated cryptic exon (CE) expression were sufficient to distinguish molecular subgroups. Interestingly, one-vs-rest subgroup discrimination based on TDP-43 associated cryptic exon (CE) expression was next assessed (new Table 3). For S2 patients, subgroup membership could be identified with near-perfect accuracy using a combination of four CE events: MYO18A CE, DNM1 CE, SYT7 CE, and ACTL6B CE (AUC = 0.993). For subgroups S3 and C, a single gene expression or mRNA splicing event was sufficient to achieve perfect separation from the remaining samples (AUC = 1.0). These events were STMN2 for S3 and HDGFL2 CE for group C. In contrast, S1 showed lower discrimination performance (AUC ≈ 0.67). Although STMN2 was the most informative feature for this subgroup, no small subset of CE events achieved robust discrimination.

